# Activation of trace amine-associated receptor 1 (TAAR1) transiently reduces alcohol drinking in socially housed mice

**DOI:** 10.1101/2022.06.02.494484

**Authors:** Bartosz Adam Frycz, Klaudia Nowicka, Anna Konopka, Marius Christian Hoener, Ewa Bulska, Leszek Kaczmarek, Marzena Stefaniuk

## Abstract

Alcohol dependence is characterized by the abnormal release of dopamine in the brain reward-related areas. Trace amine-associated receptor 1 (TAAR1) is a G Protein-Coupled Receptor that negatively regulates dopamine neurotransmission and thus is a promising target in the treatment of drug addiction. However, the role of TAAR1 in the regulation of alcohol abuse remains understudied. Here, we assessed the effect of TAAR1 activation on alcohol-drinking behaviors of C57Bl/6J female mice housed in IntelliCages. The animals were administered with either vehicle or TAAR1 full selective agonist, RO5256390, and tested for alcohol consumption, alcohol preference, and motivation for alcohol seeking. We found that mice with the highest preference for alcohol (high drinkers) in the RO5256390 group consumed less alcohol and had lower alcohol preference in comparison to high drinkers in the vehicle group, during 20 h of free alcohol access (FAA). We also found decreased alcohol consumption and alcohol preference comparing all animals in the RO5256390 to all animals in the vehicle group, during 20 h of FAA performed after the abstinence. These effects of RO5256390 lasted for the first 24 hours after administration that roughly corresponded to the compound level in the brain, measured by mass spectrometry. Finally, we found that administration of RO5256390 may attenuate motivation for alcohol seeking. Taken together, our findings reveal that activation of TAAR1 may transiently reduce alcohol drinking; thus, TAAR1 is a promising target for the treatment of alcohol abuse and relapse.

## 1. INTRODUCTION

Alcohol is one of the most abused drugs.^1^ Chronic use of alcohol can lead to an increased risk of health problems and the development of addiction referred to as alcoholism. The development of alcoholism is associated with physiological changes in the brain’s neuronal pathways that control stress and reward behaviors.^2^ One of the most affected by both binge and chronic alcohol consumption is the mesocorticolimbic system, which comprises dopamine neurons projecting from the ventral tegmental area (VTA) to multiple downstream structures, including the nucleus accumbens (NAcc), prefrontal cortex, and the amygdala. Dopamine neurons of the mesocorticolimbic pathways are essential mediators in associative learning, natural reward like food, and reinforcement of drugs of abuse.^3^ Direct administration of ethanol into the VTA of alcohol-preferring rats promoted its reinforcing effects.^4,5^ Furthermore, acute ethanol increased the release of dopamine in the NAcc^6^ and the prefrontal cortex^7^ in these animals. Thus, alcohol-induced neuroadaptation in the mesocorticolimbic system can underlie the reinforcement effect of this drug.^8,9^

In 2001, two independent research groups identified trace amine-associated receptor 1 (TAAR1) which has been further demonstrated to be involved in a broad range of neuropsychiatric disorders, including drug addiction.^10–12^ TAAR1 is an intracellular G_s_- and G_q_ Protein-Coupled Receptor (GPCR) activated by an endogenous group of so-called “trace amines”, particularly by p-tyramine, β-phenylethylamine, octopamine as well as by certain psychoactive drugs, i.e., amphetamine and methamphetamine.^13^ In the mammalian brain, TAAR1 expression was detected in the dopaminergic (VTA, substantia nigra, SN, dorsal and ventral striatum), glutamatergic (frontal cortex, amygdala, subiculum), and serotonergic (dorsal raphe) neurons.^10,14^ The study shows that TAAR1 inhibits dopaminergic transmission through modulation of the dopamine transporter (DAT)^15^, heterodimerization with dopamine receptor 2^16^, and/or activation of inwardly rectifying K^+^ channels.^17^ Furthermore, TAAR1 agonists have been shown to attenuate drug-induced dopamine secretions in structures of the mesocorticolimbic system, such as the NAcc.^18,19^

Although TAAR1 may inhibit compulsive behaviors of some drugs of abuse^20–22^, the role of TAAR1 in the regulation of alcohol-drinking behaviors is poorly understood. Recent studies have shown that TAAR1 KO mice had a higher preference and consumed more alcohol in comparison to WT mice.^23^ Furthermore, the selective partial agonist of TAAR1, RO5263397, significantly decreased the expression of ethanol-induced behavioral sensitization both in male and female WT mice.^24^

In the present study, we aimed to find out whether the intraperitoneal (IP) administration of a full selective agonist of TAAR1, RO5256390, affects the alcohol-drinking behaviors of mice housed in IntelliCage. We also studied the blood-brain barrier penetration of RO5256390 by mass spectrometry.

Similarly, to the previous reports^25^ we showed a high penetration rate of the blood-brain barrier by RO5256390 through IP administration. The IP injection of the RO5256390 reduced alcohol consumption and alcohol preference in mice with the highest alcohol preference (high drinkers) in comparison to high drinkers in the vehicle group, during the 20 h of free alcohol access (FAA). Furthermore, administration of RO5256390 decreased alcohol consumption and alcohol preference during 20 h FAA performed after abstinence, comparing all animals in the RO5256390 and the vehicle group. These effects lasted for 24 h and corresponded to the compound brain concentration. Finally, we found that mice treated with RO5256390 achieved a lower score in the motivation test in comparison to the motivation test performed before the treatment.

These results suggest that administration of RO5256390 decreases alcohol-drinking behaviors in mice, and thus TAAR1 can be a promising target in novel therapies for alcoholism.

## 2. METHODS AND MATERIALS

### 2.1 Mice

The experiments were carried out on adult 8-week C57Bl/6J female mice (n=43), bred at the animal house of the Medical University of Białystok. Animals were housed in IntelliCages either 15 or 13 mice per cage, in rooms with a controlled temperature of 22°C under a 12/12 h light-dark cycle. The cage environment was enriched with shelters, nesting material, and gnawing sticks. Before caging, mice were anesthetized with isoflurane and were implanted with RFID transponders (8.5 mm in length and 1.4 mm in diameter). Mice were weighed three times during the study. After behavioral studies, mice were sacrificed by decapitation under isoflurane anesthesia. All procedures were performed following the Animal Protection Act in Poland, Directive 2010/63/EU, and were approved by the 1st Local Ethics Committee (Permission no. 793/2018).

### 2.2 IntelliCages

Mice were tested in three automated learning systems (IntelliCages) from TSE systems (Berlin, Germany) [www.tse-systems.com].^26,27^ Two corners of the cage were equipped with a presence detector and an antenna to detect the signal from the RFIDs. Access to bottles with water and/or alcohol was limited by a door behind the entrance of the corner. Activity towards the entrance (nose-poke) opened the door and allowed the mouse to lick from the bottle (Figure 1A). Visits to the corner with alcohol were followed by a green led light. The activity of mice in a corner, including visits, nose-pokes, and licks was automatically recorded. Preference for alcohol drinking, alcohol consumption, and motivation for alcohol seeking was examined by adequate cage setup (Figure 1B).

**FIGURE 1.**
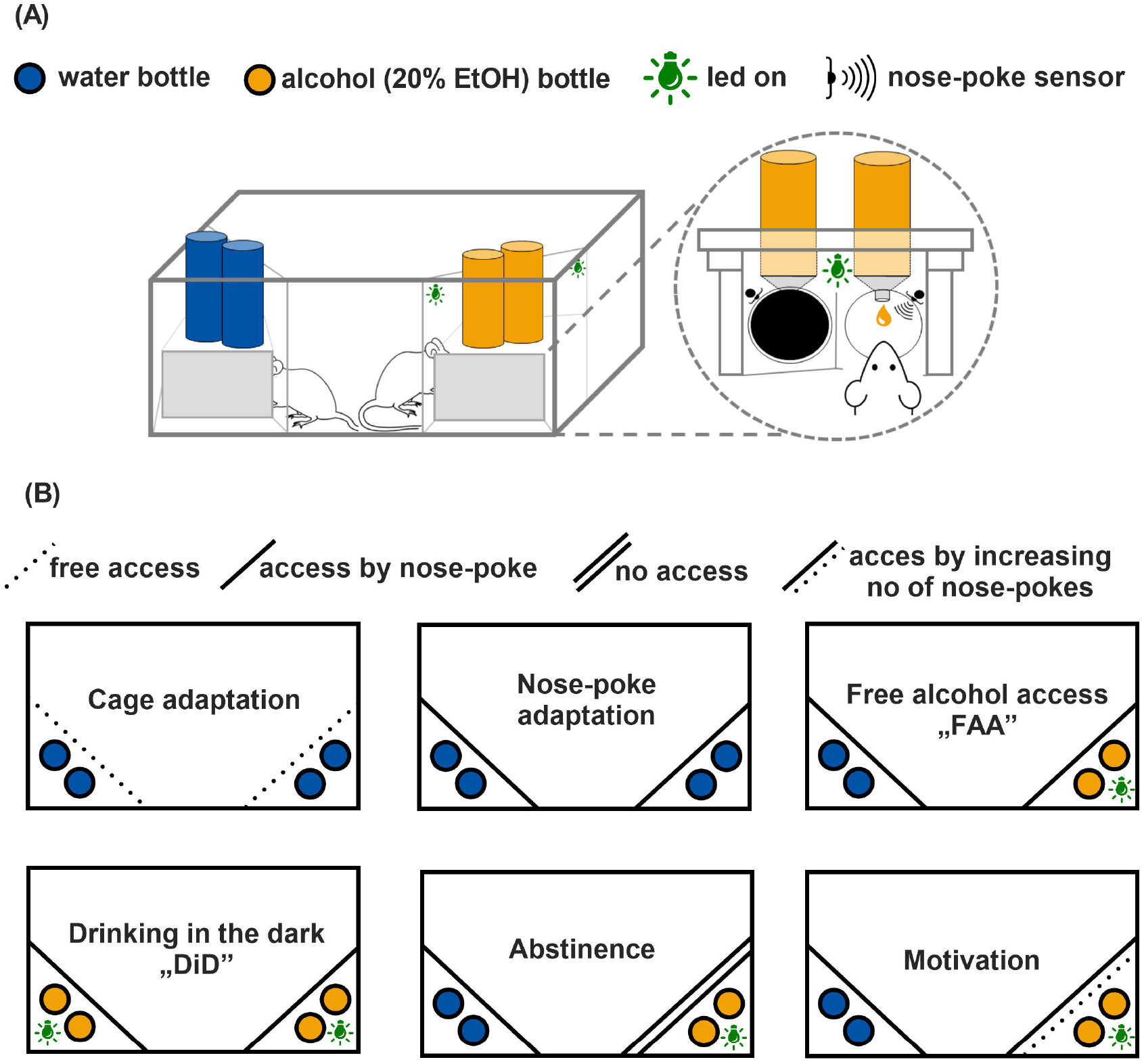
IntelliCage was used to study the effects of RO5256390 administration on alcohol drinking in C57Bl/6J mice. (A) To access water or alcohol, a mouse had to enter the corresponding corner and approach a sensor (nose-poke), which opens a door blocking access to the bottle. (B) During the adaptation and free access to alcohol (FAA), mice had unrestricted access to water or water and alcohol, respectively. The FAAs were preceded by a Drinking in the Dark (DiD) that involved replacing all water bottles with bottles containing 20% ethanol [v/v] for 2.5 h, beginning 2 h into the dark cycle. During the motivation tests, the animals had to perform an increasing number of nose-poke at each visit to get access to alcohol. In the abstinence period, access to alcohol was blocked. The presence of mice in a corner with alcohol was always accompanied by the lighting of green led.

### 2.3 Behavioral training

Mice were placed in IntelliCages and underwent place adaptation for 10 days. Then, they underwent behavioral training consisting of a series of sessions based on the drinking in the dark paradigm (DiDs)^28^ and sessions with free access to alcohol (FAAs). Briefly, DiDs involved replacing all water bottles with bottles containing 20% ethanol [v/v] for 2.5 h, beginning 2 h into the dark cycle. The presence of mice in corners with alcohol was accompanied by the lighting up of green light. DiDs lasted 4 weeks, every day, excluding one day in a week when mice only had access to water. DiDs was then followed by the 3 weeks of FAAs. FAAs were conducted similarly to DiDs, except that half of the bottles contained water. Thus, mice could choose whether they preferred to drink water or alcohol. Bottles of alcohol were placed in a corner that was previously visited less frequently by mice to exclude the effect of place preference. Mean consumption of alcohol in the pre-treatment DiDs and FAAs (2.5 h) as well as mean preference for alcohol drinking during FAAs were calculated. The following formula was used to calculate the consumption of alcohol: (number of licks × 1.94 × alcohol concentration × 0.79 g/ml)/animal weight. The value of 1.94 μl ± 0.2 μl corresponds to the average lick volume. Alcohol consumption was presented as g of consumed alcohol per kg of body weight (g/kg). Preference for alcohol was calculated as the total number of ethanol licks divided by the total number of all licks performed in both, the alcohol and water corners, and expressed as a percentage.

### 2.4 Experimental groups

Mice (n=30) were assigned to the “vehicle” and “RO5256390” experimental groups. Similarly, to Radwanska and Kaczmarek (2011)^29^ the 30% of animals in each group that had the highest alcohol consumption and alcohol preference during behavioral training were classified as “high drinkers” while the other animals were classified as “low drinkers” (Supporting Table). There was 5 high, and 10 low drinkers in each group. Average alcohol consumption and preference for alcohol were similar in both groups before treatment (Supporting Table). Motivation for alcohol seeking and the long-drinking assay was performed on another group of mice (n=13) assigned to the vehicle or RO5256390 group based on their result in motivation tests before treatment. The average motivation score between these groups obtained during motivation tests performed before treatment was similar (Figure 5B).

**FIGURE 2.**
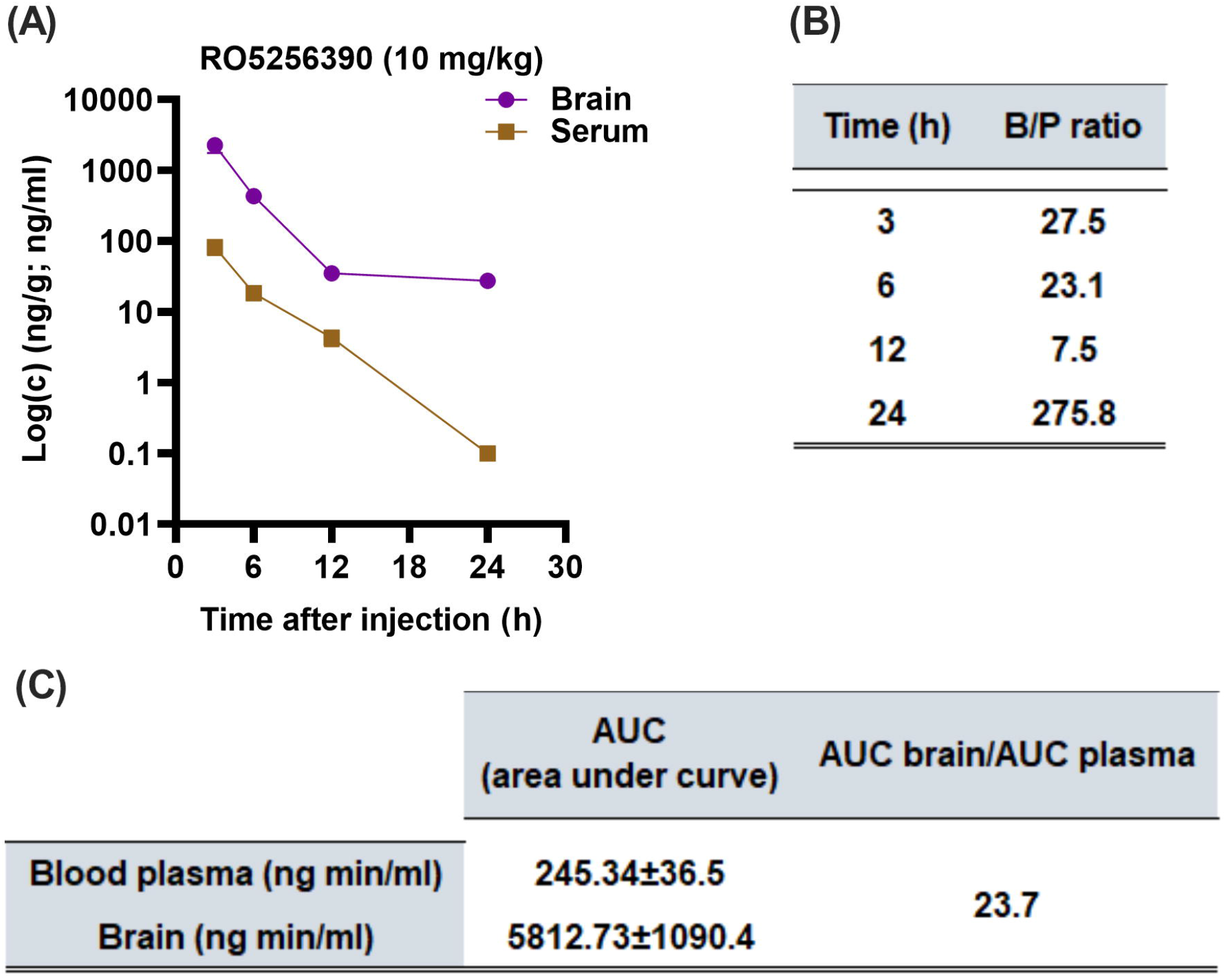
RO5256390 effectively penetrated the blood-brain barrier. (A) RO5256390 was found in both, the brain and plasma after IP administration. (B) RO5256390 showed a brain/plasma ratio (B/P) >1 in all time points which indicates a high blood-brain barrier penetration rate of the compound. (C) The high penetration rate of the blood-brain barrier by RO5256390 was additionally demonstrated by the area under the curve (AUC) calculations.

**FIGURE 3.**
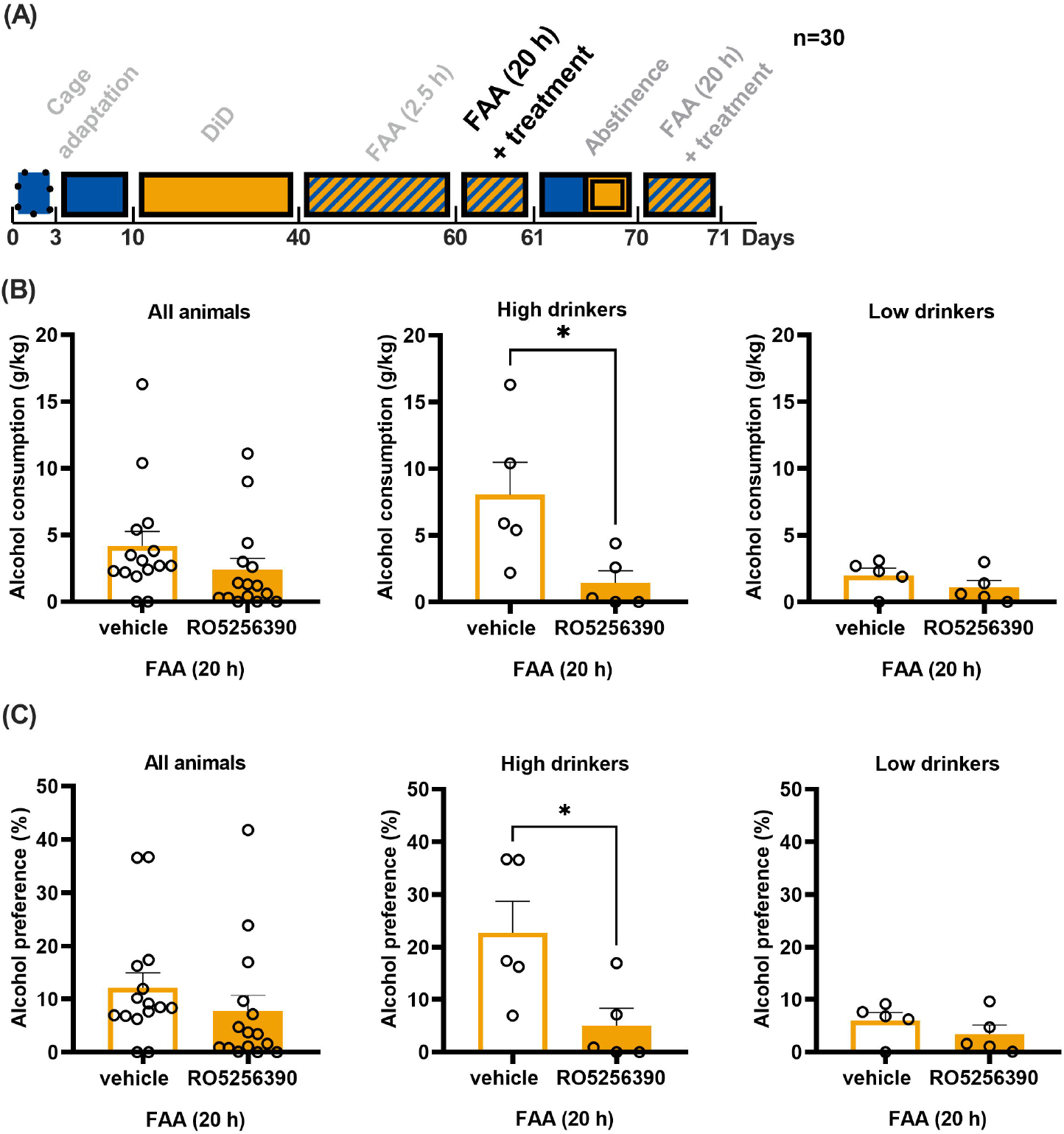
RO5256390 administration reduced alcohol consumption and alcohol preference in high drinkers during free alcohol access (FAA). (A) Experimental timeline. (B) Total alcohol consumption during FAA was lower in high drinkers in the RO5256390 group in comparison to high drinkers in the vehicle group (p=0.035, Unpaired t-test). (C) RO5256390 decreased alcohol preference in high drinkers in comparison to the high drinkers in the vehicle group (p=0.031, Unpaired t-test).

**FIGURE 4.**
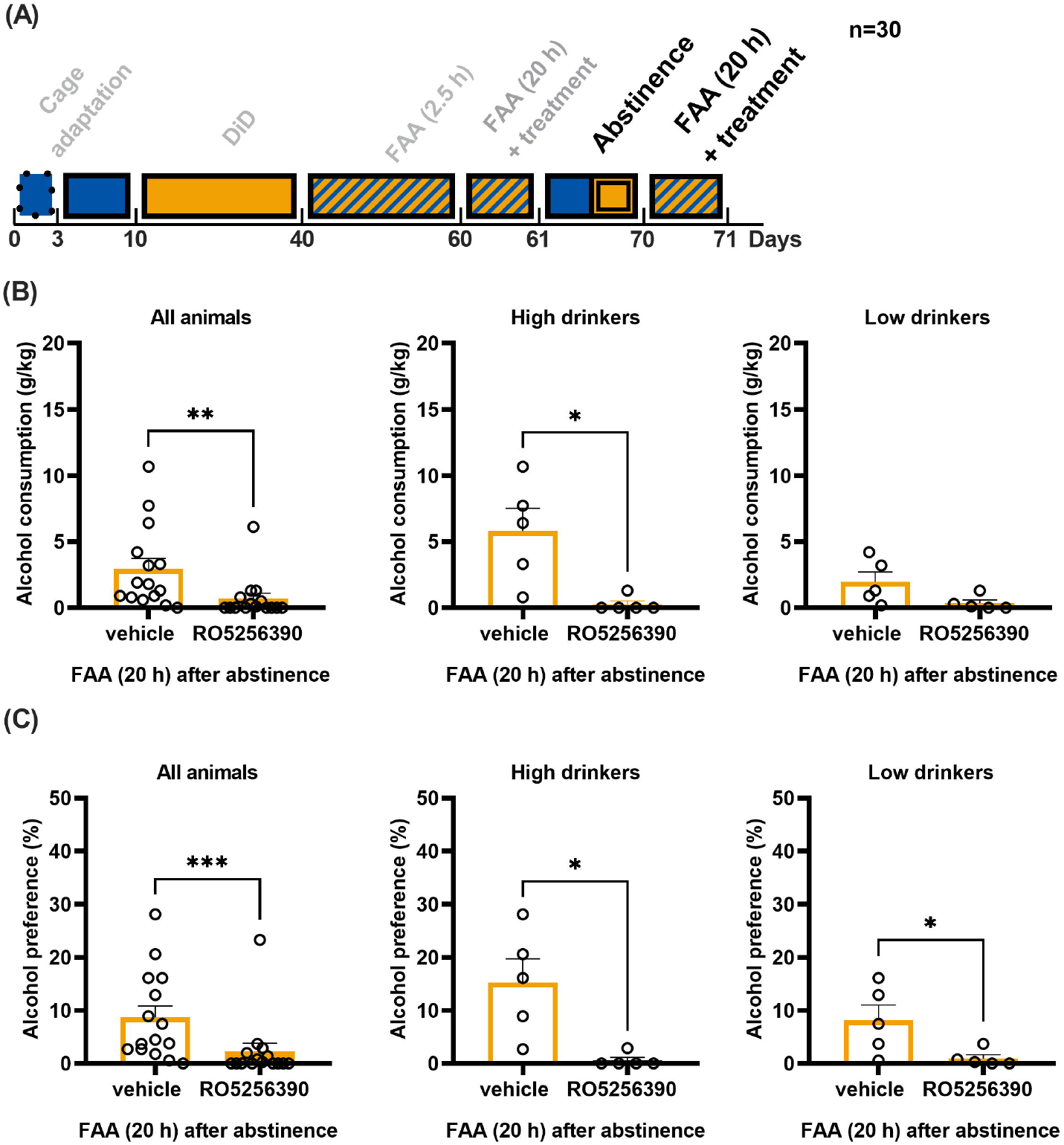
RO5256390 administration reduced alcohol consumption and alcohol preference during free alcohol access (FAA) performed after abstinence. (A) Experimental timeline. (B) Total alcohol consumption during FAA performed after abstinence was lower comparing all animals in the RO5256390 group to their counterparts in the vehicle group (p=0.0011, Mann-Whitney U test). (C) RO5256390 decreased alcohol preference comparing all animals in the RO5256390 group to their counterparts in the vehicle group (p=0.0009, Mann-Whitney U test).

**FIGURE 5.**
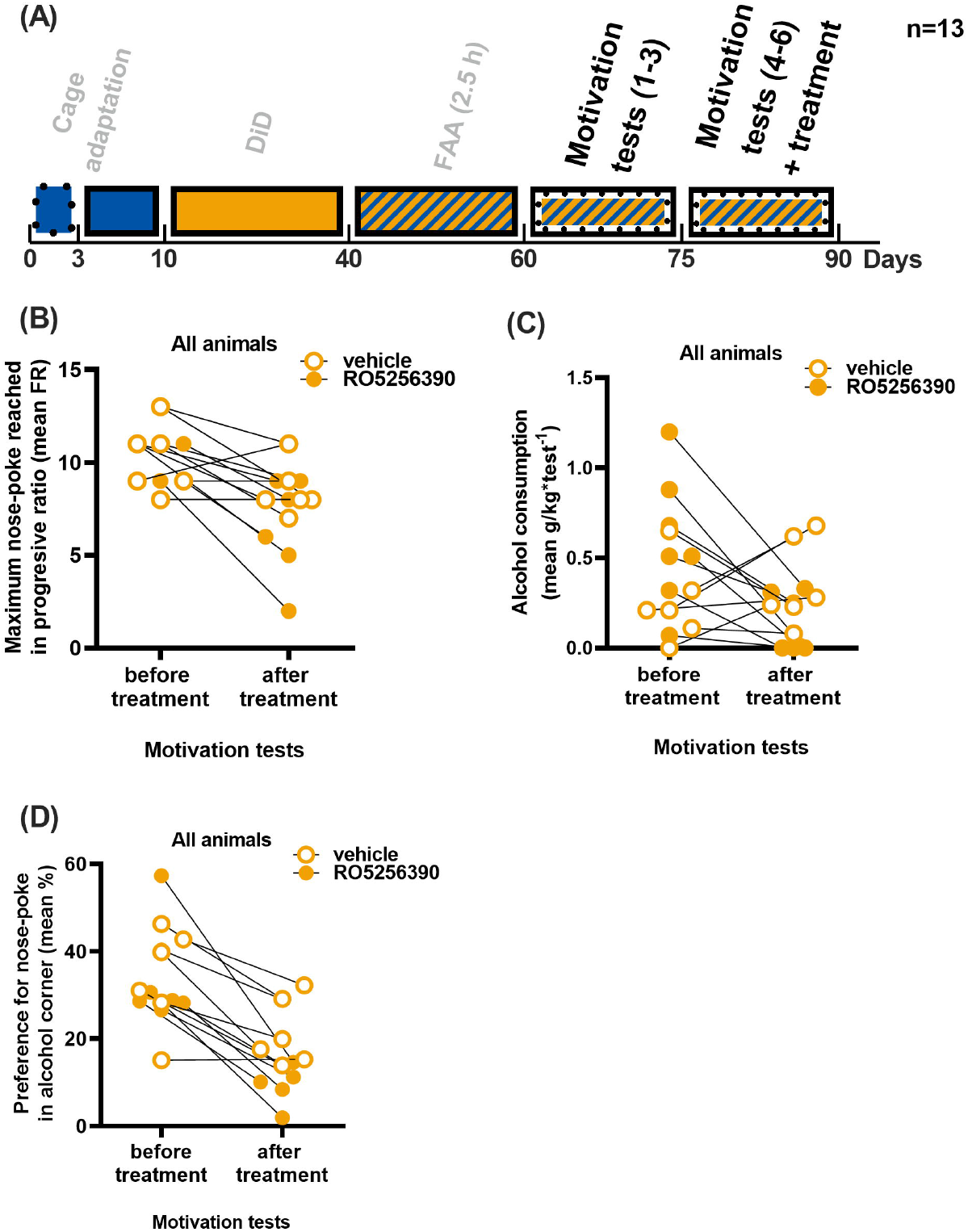
RO5256390 administration attenuated motivation for alcohol drinking. (A) Experimental timeline. (B) RO5256390 mice but not vehicle group mice achieved a statistically significant lower score in the motivation tests after compound administration in comparison to the pre-treatment tests score (p=0.018, Tukey’s multiple comparisons). (C) Mice treated with RO5256390 consumed less alcohol (p=0.015, Tukey’s multiple comparisons) in motivation tests performed after the treatment in comparison to the consumption in the motivation tests performed before the treatment (D) Mice in RO5256390 group had decreased preference (p=0.0023, Tukey’s multiple comparisons) for doing nose-pokes in the alcohol corner during post-treatment tests in comparison to pre-treatment tests.

### 2.5 Drugs

The highly selective TAAR1 full agonist, RO5256390, and Kolliphor EL were purchased in Merck/Millipore Sigma (Burlington, MA, United States). RO5256390 was dissolved in a mixture of ethanol (5% v/v), Kolliphor EL (10% v/v), and saline, just before the start of the experiment. Both RO5256390 and the vehicle were injected intraperitoneally (IP). The dose was expressed as an mg of the compound per kg of body weight (mg/kg). The dose of 10 mg/kg for RO5256390 was based on previously published reports.^30,31^

### 2.6 RO5256390 administration and behavioral tests

Mice were habituated for IP injection by the tail and scruff handling and mimicking injection, for 10 days before the treatment. Mice (n=30) then were treated with either vehicle or RO5256390 followed by 20 h FAA. During the treatment, FAAs were extended to 20 h instead of 2.5 h due to the low activity of RO5256390-treated mice in corners during the first 2 h after drug administration (data not shown). The effect of RO5256390 on alcohol drinking after abstinence was investigated in a 20 h FAA performed after blocking access to the alcohol corner for 9 days. The study of the effect of RO5256390 on motivation for alcohol-seeking was investigated in another group of mice (n=13). Like previously, mice underwent behavioral training consisting of DiDs and FAAs. Then, they underwent 6 motivation tests that were conducted at an interval of one week. The score in motivation tests was represented by the maximum number of nose-poke a mouse performed to drink alcohol during a single visit to the corner. The number of nose-pokes required to open the door to alcohol increased after each successful attempt, according to the following ratio sequence (FR): 1, 4, 8, 12, 16, 20, 24, 28, and 32. Each test lasted 20 h. Mice were habituated to IP as previously. The same animals were subsequently tested for the effect of RO5256390 on long-term alcohol drinking that included administration with either vehicle or RO5256390 and 48 h FAA.

### 2.7 Blood-brain barrier penetration

#### 2.7.1 Extraction procedure

Mice (n=12) were injected with RO5256390 at the dose of 10 mg/kg. The animals were sacrificed at 3, 6, 12, and 24 h post-injection, and blood and brain tissue samples were collected. Three animals were sacrificed for every time point. The mice were perfused with 0.9% NaCl before the brain collection. Blood was subjected to centrifugation (2000 x g for 10 min) in the presence of sodium citrate to obtain a clear plasma sample. Brain tissues were weighted using an analytical scale and homogenized in 1 ml of deionized water using TissueRuptor II electric homogenizer by Qiagen (Germantown, MA, USA). RO5256390 was extracted using acetonitrile/water 50%/50% (v/v) (LC/MS grade, JT Baker) as an extraction solution. Briefly, 100 μl and 50 μl of brain homogenate and plasma samples were added to 500 μl and 250 μl of extraction buffer, respectively. Samples were incubated for 10 min. in a thermomixer (Eppendorf) at room temperature at 800 × rpm. Supernatants were separated from residues by centrifugation for 10 min. at 17 500 × g (Centrifuge 5804/5804R, Eppendorf). Pellets were discarded, whereas 200 μl of supernatants were transferred to LC autosampler vials (Agilent Technologies) and analyzed by UHPLC-ESI-MS/MS. The extraction procedure was carried out in three technical replicates for each examined tissue sample.

#### 2.7.2 UHPLC-ESI-MS/MS analysis

UHPLC-ESI-MS/MS analysis was performed using an Agilent 1290 Infinity LC system (Agilent Technologies) coupled to Agilent 6460 Triple Quadrupole Mass Spectrometer (Agilent Technologies) equipped with an electrospray ionization source (ESI) operating in positive ion mode. ESI conditions were as follows: nebulizing gas temperature – 300 °C, nebulizing gas flow – 10 l/min., sheath gas temperature – 200 °C, sheath gas flow – 11 l/min., positive ionization mode source voltage – 3.75 kV. Nitrogen was used as nebulizing and sheath gas. The mass spectrometer was operated in the multiple reaction monitoring mode (MRM) following optimization of working conditions (fragmentor voltage and collision energy, CE) for RO5256390 using a standard solution at the concentration of 1 mg/L (in acetonitrile/water 50%/50% (v/v) solution) in the direct infusion mode. The optimal fragmentor voltage was found to be 80 V. The following transitions of m/z (precursor ion → product ion) were monitored: 219.1 → 159.1 (CE=8 eV) (as the most intense transition taken for quantitative analysis) and 219.1 → 117.1 (CE=19 eV), 219.1 → 91.1 (CE=41 eV), 219.1 → 61.2 (CE = 12 eV) (confirming transitions for qualitative analysis). Compounds, present in the prepared extracts, were separated on a Zorbax Eclipse Plus C18 Rapid Resolution column (Agilent Technologies) (100 mm x 4.6 mm, 3.5 μm) at 35 °C. Water with 0.1% formic acid and acetonitrile with 0.1% formic acid was used as eluent A and B, respectively. The mobile phase was delivered at 0.5 mL/min in isocratic mode with 50% of eluent B. The injection volume was 10 μl. The total time of chromatographic separation was 10 min. Measurements were carried out in three technical replicates. Quantification of RO5256390 was achieved by the external matrix calibration curve method. Standard solutions were made by adding appropriate amounts of RO5256390 stock solution to extracts of blank brain homogenates and plasma. Blank extracts were obtained from tissues derived from untreated animals following the same extraction procedure. Standard curves for brain aliquots and blood plasma were generated within the concentration range of 0.1 ng/ml – 100 ng/ml (R^2^=0.9999) and 0.1 ng/ml – 50 ng/ml (R^2^=0.9995), respectively. Considering the dilution factor and the masses of tissues collected, the content of RO5256390 was calculated per g of tissue or ml of plasma for brain tissue and blood plasma samples, respectively.

### 2.8 Statistical analysis

All results are presented as mean ± SEM. The appropriate tests were used, considering whether the data had normal distribution and equal variance. Differences in alcohol consumption between mice in the RO5256390 and vehicle groups were estimated using the Unpaired t-test or Mann-Whitney U test. The effect of the drug administration on motivation for alcohol seeking was evaluated by Tukey’s multiple comparisons test. Statistical analysis was conducted using GraphPad Prism, version 9.3.1 (GraphPad Software, Inc., La Jolla, CA).

## 3. RESULTS

### 3.1 RO5256390 highly penetrates the blood-brain barrier

RO5256390 was detected in both blood plasma and the brain after IP administration (Figure 2A-C). The highest concentration of RO5256390 in the brain and plasma was observed 3 h post-injection (2260 ng/g_(*brain*)_ and 82ng/ml_(*plasma*)_, respectively). The amount of RO5256390 in these tissues was decreasing over time. Twenty-four hours after injection, the amount of RO5256390 in plasma was at the trace level (<0.1 ng/ml) while in the brain it was 27.6 ng/g (1.2% of the maximum amount reached at 3 h post-injection) (Figure 2A, B). Notably, the B/P ratios for all studied time points exceeded 1, demonstrating higher brain RO5256390 concentrations than plasma concentrations (Figure 2B). Furthermore, we calculated the ratio of areas under the kinetic curve (AUC brain/AUC plasma) of the RO5256390 concentration in blood plasma and brain tissue (Figure 2C). The ratio of 23.7 shows the high ability of RO5256390 to penetrate the blood-brain barrier.

### 3.1 Mice showed different alcohol preferences during behavioral training

We used a model of binge-like ethanol drinking, “drinking in the dark” (DID) as well as free alcohol access (FAA) to select the animals with the highest preference for alcohol. Average alcohol consumption during 2.5 h DiDs was 2.28 g/kg (range: 1.20 - 3.88 g/kg; high drinkers vs. low drinkers: 2.75 vs. 2.04 g/kg, Supporting Table). Average alcohol consumption during 2.5 h FAAs was 0.71 g/kg (range: 0.00 – 1.57 g/kg; high drinkers vs. low drinkers: 1.20 vs. 0.46 g/kg, Supporting Table). Average alcohol preference during 2.5 h FAAs was 27.25% (range: 0.00 – 64.79%; high drinkers vs. low drinkers: 38.23% vs. 21.76%, Supporting Table)

### 3.2 RO5256390 decreases alcohol consumption and alcohol preference

Mice were treated with either RO5256390 or the vehicle and tested for alcohol consumption and alcohol preference (Figure 3A-C). We found that high drinkers treated with RO5256390 consumed less alcohol (p=0.035) (Figure 3B) and had decreased preference for alcohol (p=0.031) (Figure 3C) in comparison to high drinkers treated with vehicle, during 20 h of FAA. Simultaneously, there were no differences in water consumption between these groups (Supporting Figure). Then, we studied the effect of RO5256390 administration on alcohol drinking and alcohol preference after the abstinence period (Figure 4A-C). We found decreased alcohol consumption (p=0.0011) (Figure 4B) and alcohol preference (p=0.0009) (Figure 4C) comparing all animals in RO5256390 to their counterparts in the vehicle group. The water consumption during the test was similar in both groups (Supporting Figure).

### 3.3 RO5256390 attenuates motivation to alcohol drinking

We compared the pre-, and post-treatment results obtained in motivation tests by mice in vehicle and RO5256390 groups (Figure 5A-D). We found that RO5256390-treated animals scored lower (p=0.018) on post-treatment motivation tests in comparison to the pretreatment score (Figure 5B). Mice in the RO5256390 group also consumed less alcohol (p=0.015) in motivation tests performed after the treatment in comparison to the consumption in the motivation tests performed before the treatment (Figure 5C). Furthermore, mice treated with RO5256390 had decreased preference (p=0.0041) for doing nose-pokes in the alcohol corner during post-treatment tests in comparison to pre-treatment tests (Figure 5D).

### 3.4 The effect of RO5256390 on alcohol drinking is short-lasting

We studied the duration of reduced alcohol drinking in mice treated with RO5256390 (Figure 6A-C). We observed reduced alcohol consumption only during the first 24 h after the treatment (p=0.024) (Figure 6B) whereas water consumption was similar for both groups throughout the entire experiment (Figure 6C). These results are consistent with our previous experiments where the RO5256390 group consumed a similar amount of alcohol in comparison to the vehicle group in FAA performed the next days after treatment (data not shown).

**FIGURE 6.**
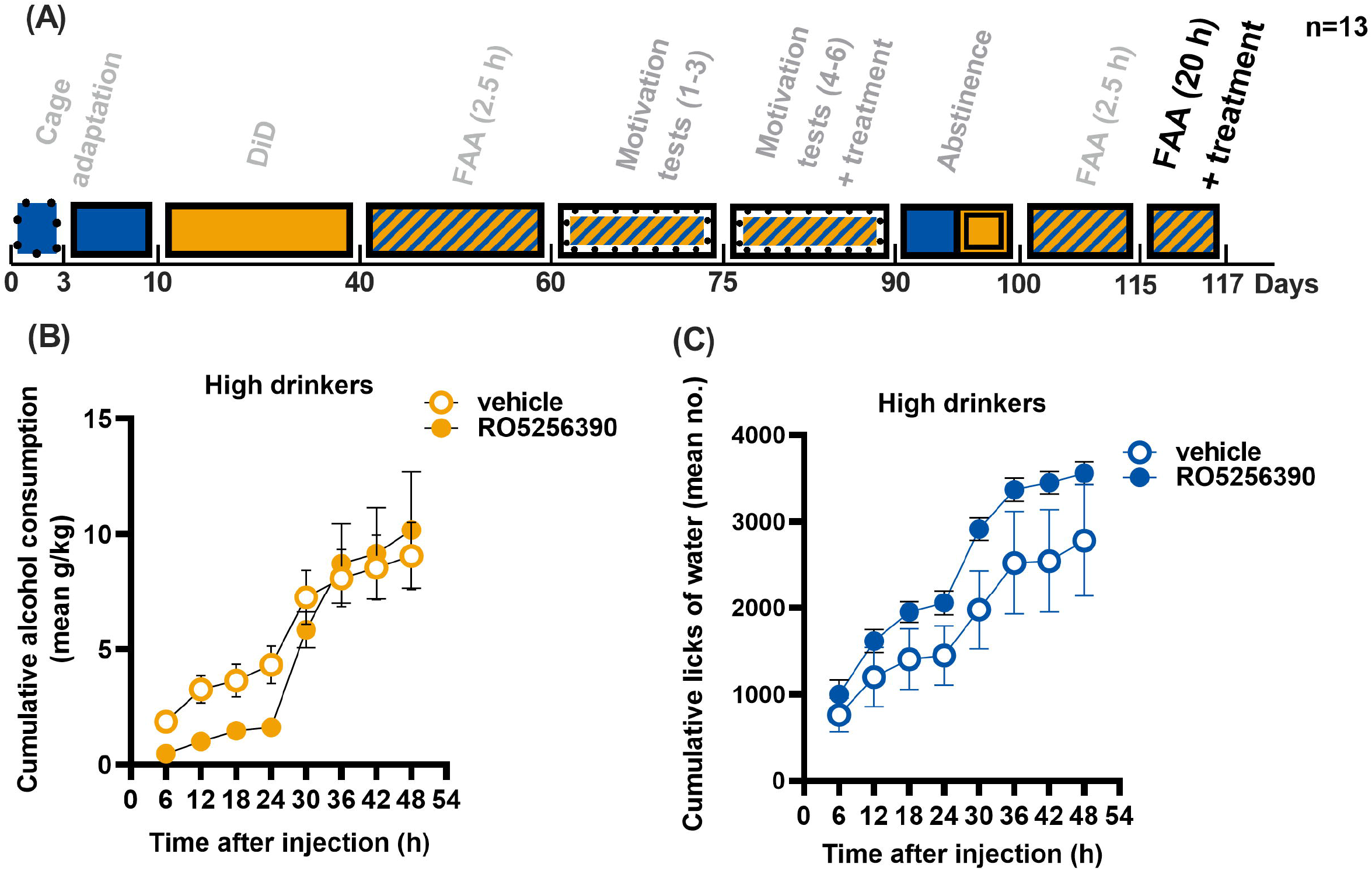
Reduced alcohol consumption after RO5256390 administration is short lasting. (A) Experimental timeline. (B) High drinkers in the RO5256390 group consumed less alcohol 24 h after compound administration in comparison to the high drinkers in the vehicle group (p=0.024, Unpaired t-test). The difference was lost during the following 24 h. (C) There were no differences in the number of licks in the water corner between RO5256390 and vehicle groups within 48 h free alcohol access.

## 4. DISCUSSION

Our study reports that the full selective TAAR1 agonist, RO5256390, reduces alcohol drinking in female C57BL/6J mice compared to their littermates administered with a vehicle. Mice in the RO5256390 group also consumed less alcohol in comparison to vehicle mice after the abstinence period. Furthermore, we have shown that RO526390 administration can attenuate motivation for alcohol seeking. We also confirmed that RO5256390 effectively penetrates the blood-brain barrier after IP administration.

Our results suggest that activation of TAAR1 decreases alcohol drinking in mice which appears to be consistent with the previous studies. Lynch et al.^23^ showed that TAAR1 deletion in mice significantly increases their alcohol consumption and alcohol preference in comparison to WT counterparts. This suggests the role of TAAR1 in the regulation of alcohol drinking.

It must be emphasized that we analyzed voluntary alcohol drinking by a group of mice socially housed in environmentally enriched cages, and the animals here were not artificially motivated to drink the ethanol solution (except for the 2.5 h of water deprivation during the DiDs). Holgate et al.^32^ demonstrated that mice housed socially in IntelliCage with environmental enrichment had even ~8 times less preference for alcohol than mice housed individually (8.35 ± 1.570% vs. 65.20 ± 1.747%). Alcohol consumption of mice in our study was up to about 3 times lower than that of mice housed individually elsewhere.^33^ Therefore, due to the likely low blood alcohol concentration (BEC) obtained in our approach, the model we used is not a model of addiction. Rather it represents alcohol drinking under an intermittent access schedule in group-housed mice. The main advantages of this approach are the reproducibility of the alcohol preference and testing animals under their social interactions.^34,35^ Lower consumption of alcohol by mice in our study may also be caused by not using sucrose to motivate animals to drink. Although TAAR1 seems not to affect natural reward ^23,36^, in our approach we wanted to exclude the potential effect of sucrose on alcohol consumption. Importantly, it was shown that environmental enrichment may impacts differently on alcohol than sucrose consumption^32^; thus, sucrose may not be suitable for our studies.

Wu et al.^24^ showed decreased expression of ethanol-induced behavioral sensitization both in male and female WT mice treated with a highly selective partial TAAR1 agonist, RO5263397. Since there is a remarkable overlap in the neurocircuitry between behavioral sensitization and relapse, TAAR1 activation may affect these phenomena. Consistently, we found a strong effect of RO5256390 on alcohol drinking performed after the abstinence period, which in our study mimics the psychological discomfort of prolonged alcohol deprivation. Similar effects were observed in the psychostimulant study. Administration of RO5256390 produced a downward shift in the dose-response curve for cocaine self-administration and dose-dependently prevented the cocaine-induced lowering of intracranial self-stimulation thresholds^37^, and dose-dependently decreased cocaine seeking in rats, after a 2-week withdrawal from chronic cocaine self-administration.^30^ Interestingly, recent studies have shown that RO5256390 can regulate compulsive behavior related to food intake. It was found that the compound dose-dependently blocks binge-like eating of a palatable diet in rats.^38^ RO5263397 also blocked methamphetamine-primed reinstatement of the drug-seeking which was accompanied by a decrease in the release of dopamine in NAcc.^19^ These suggest that activation of TAAR1 may attenuate the rewarding effects of the drugs or binge eating, and prevent relapse-like behaviors.

We do not know the mechanism by which TAAR1 activation resulted in reduced alcohol consumption in mice from our study. It cannot be excluded that the effects we observed are related to RO5256390 impact on the taste of alcohol. Although selective agonists of TAAR1 attenuated addiction-related behaviors without producing aversive effects themselves, there may be potential aversive effects elicited by these drugs.^39^ Several studies showed that trace amines, including tryptamine and phenethylamine, produce taste aversion to saccharin novelty.^40–42^ Furthermore, significant aversions to both saccharin and NaCl taste novelty were produced by RO5166017 and RO5263397.^43^ Harkness et al.^44^ also showed that stimulation of a sub-functional TAAR1 may increase sensitivity to the aversive effects of methamphetamine. Therefore, TAAR1 activity may increases sensitivity to the aversive effects of the drug, including alcohol.

Similarly to the previous studies,^29^ we adapted a progressive-ratio test of mice’s motivation for alcohol seeking. We found that the RO5256390 administration attenuated the motivation for alcohol seeking. Similarly to this observation, RO5263397 decreased the motivation for methamphetamine self-administration under a progressive ratio schedule of reinforcement.^19^ The same compound also significantly decreased nicotine’s physical and motivational withdrawal effects in the long-access self-administration rats model.^45^ This indicates the possible involvement of TAAR1 activation in the regulation of motivational behavior associated with drug addiction.

Although it has been quite well established that alcohol consumption differs between male and female C57Bl/6 mice,^46^ we were unable to study gender effect in RO5256390 treatment. IntelliCage has been used successfully in the long-term behavioral studies of female mice, while males would eventually require additional compartment barriers.^46^ To limit any potential dominance fights that could disturb the results we decided to only use female mice. On the other hand, evidence suggests that women develop an addiction to alcohol much more quickly than men.^47^ Consistently, female mice had augmented ethanol-induced locomotor activity in comparison to male individuals.^24^ There is also evidence of increased alcohol consumption in female than male rats, occurring after the second and third deprivations.^48^ However, it remains unknown whether the administration of TAAR1 agonists affects differently on males and females.^49^

RO5256390 has been recognized as a potent full agonist at hTAAR1 and has excellent pharmacokinetic properties in mice and rats.^50,51^ We confirmed that the compound highly penetrates the blood-brain barrier in mice. Despite the low to moderate oral bioavailability of RO5256390, Revel et al.^50^ showed that IP administration of the compound to mice increases its bioavailability. An elimination half-life of R05256390 was estimated to be several hours with a high brain/plasma concentration ratio (11 times more in the brain).^50^ In our study, 3 h after administration, the amount of RO5256390 in the brain was nearly 30 times higher than in plasma and was approx. 30% of the original dose. However, we used a higher dose than Revel et al. (10 mg/kg vs 2.1 mg/kg IP). Interestingly, 24 h after administration, the amount of RO5256390 in both brain and plasma was drastically low compared to the original dose. It is worth noting here that the reduction in alcohol intake in RO5256390 mice in comparison to vehicles resolved after approximately 24 h and remained similar in both groups thereafter.

Finally, on the second day after compound administration, we observed no differences in alcohol consumption between the studied groups. This indicates an effect of RO5256390 on drinking alcohol only while it is present in the brain. It is also important to consider that we used the whole-brain extract in mass spectrometry studies. Thus, we do not know whether RO5256390 does not better penetrate specific brain areas.

In conclusion, our findings indicate that TAAR1 activation may be involved in the regulation of alcohol-drinking-related behaviors. Further, TAAR1 agonists such as RO5256390 may be potentially used in the treatment of alcohol abuse, specifically inhibiting motivation to drink alcohol and drinking alcohol after abstinence. These all suggest that TAAR1 is a promising pharmacological target for the treatment of alcoholism; however, more research is needed.

## ACKNOWLEDGEMENTS

This work was funded by the Polish National Science Centre (NCN), Sonatina grant: 2018/28/C/NZ4/00453. The MS-based quantification analysis was performed at the Biological and Chemical Research Centre, University of Warsaw, established within the project that was co-financed by the European Union from the European Regional Development Fund under the Operational Programme Innovative Economy 2007–2013.

## CONFLICT OF INTEREST

Dr. Marius C. Hoener is employed by F. Hoffmann-La Roche AG where the RO5256390 was developed.

## AUTHOR CONTRIBUTIONS

BAF, MS, and LK were responsible for the study concept and design. BF and KN performed behavioral experiments. MCH was responsible for scientific consultation and discussion of results. AK and EB performed mass spectrometry assays. BAF wrote the manuscript. MS assisted in figure preparation.

## DATA AVAILABILITY STATEMENT

The data that support the findings of this study are available from the corresponding author upon reasonable request.

---

We found that administration of TAAR1 full selective agonist, RO5256390, transiently reduces alcohol consumption in C57BI/6J mice. Furthermore, RO5256390 may attenuate motivation for alcohol-seeking in the studied animals. Taken together, TAAR1 agonists are promising drugs for the treatment of alcohol abuse and relapse.

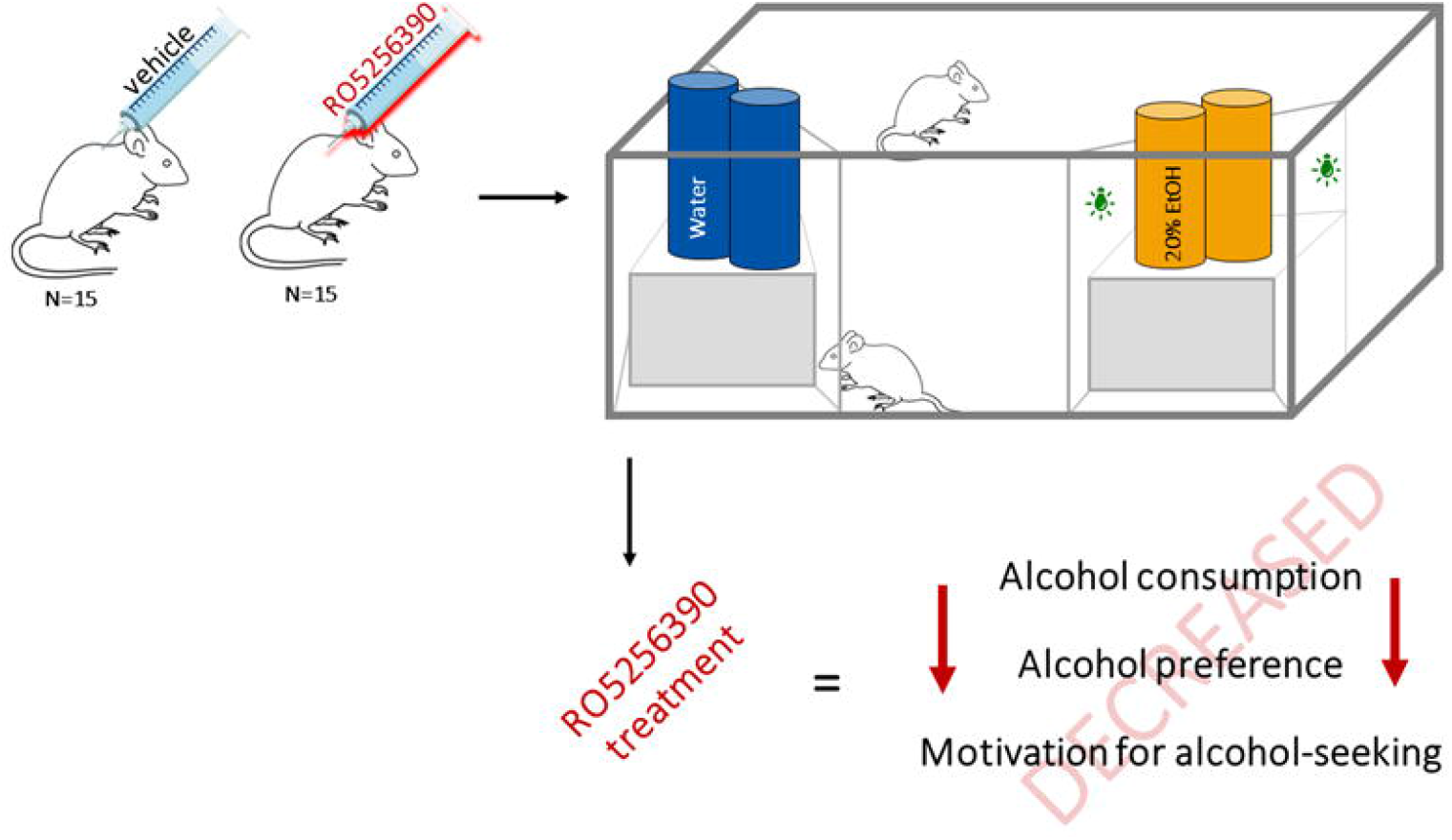

## Notes

### Summary of Updates

Discussion has been extended Figures have been modified

