## Supporting figures for "Activation of trace amine-associated receptor 1 (TAAR1) transiently reduces alcohol drinking in socially housed mice"

SUPPLEMENTARY TABLE Alcohol preference in mice before the experiments.

| **Mouse No.** | **EtOH consumption (g/kg)** | | **Alcohol preference (%)** (mean/session) |
|---|---|---|---|
| | **DiD** (mean/session) | **FAA** (mean/session) | |
| *Mice assigned to the "vehicle" group* | | | |
| **2** | 2.12 | 0.86 | 42.25 |
| **3*** | 2.06 | 1.16 | 53.30 |
| **5** | 1.64 | 0.39 | 10.48 |
| **6*** | 1.92 | 1.31 | 39.61 |
| **10** | 1.35 | 0.70 | 32.51 |
| **11** | 1.45 | 0.41 | 17.21 |
| **12** | 1.52 | 0.95 | 56.69 |
| **17** | 2.57 | 0.31 | 18.89 |
| **22** | 1.91 | 0.35 | 19.51 |
| **23** | 1.73 | 0.00 | 0.00 |
| **26*** | 3.88 | 1.25 | 37.14 |
| **27** | 2.25 | 0.36 | 14.44 |
| **28** | 3.20 | 0.68 | 22.66 |
| **29*** | 3.21 | 1.10 | 30.96 |
| **30*** | 3.35 | 1.09 | 32.48 |
| **mean** | **2.28** | **0.73** | **28.54** |
| ±SEM | 0.20 | 0.11 | 4.08 |
| *Mice assigned to the "RO5256390" group* | | | |
| **1** | 1.73 | 0.29 | 15.76 |
| **4*** | 1.75 | 1.57 | 47.18 |
| **7*** | 1.93 | 0.85 | 38.57 |
| **8** | 1.75 | 0.22 | 13.23 |
| **9** | 1.31 | 0.70 | 23.86 |
| **13** | 2.38 | 0.69 | 29.41 |
| **14** | 1.26 | 0.78 | 64.79 |
| **15** | 1.43 | 0.18 | 6.64 |
| **16** | 2.47 | 0.53 | 18.88 |
| **18*** | 3.02 | 1.13 | 51.84 |
| **19*** | 3.52 | 1.48 | 27.40 |
| **20*** | 2.82 | 1.05 | 23.81 |
| **21** | 3.81 | 0.37 | 11.36 |
| **24** | 3.75 | 0.40 | 9.85 |
| **25** | 1.20 | 0.11 | 6.75 |
| **mean** | **2.28** | **0.69** | **25.96** |
| ±SEM | 0.24 | 0.12 | 4.55 |

*high drinkers

SUPPORTING FIGURE  Water consumption of mice in free access to alcohol sessions

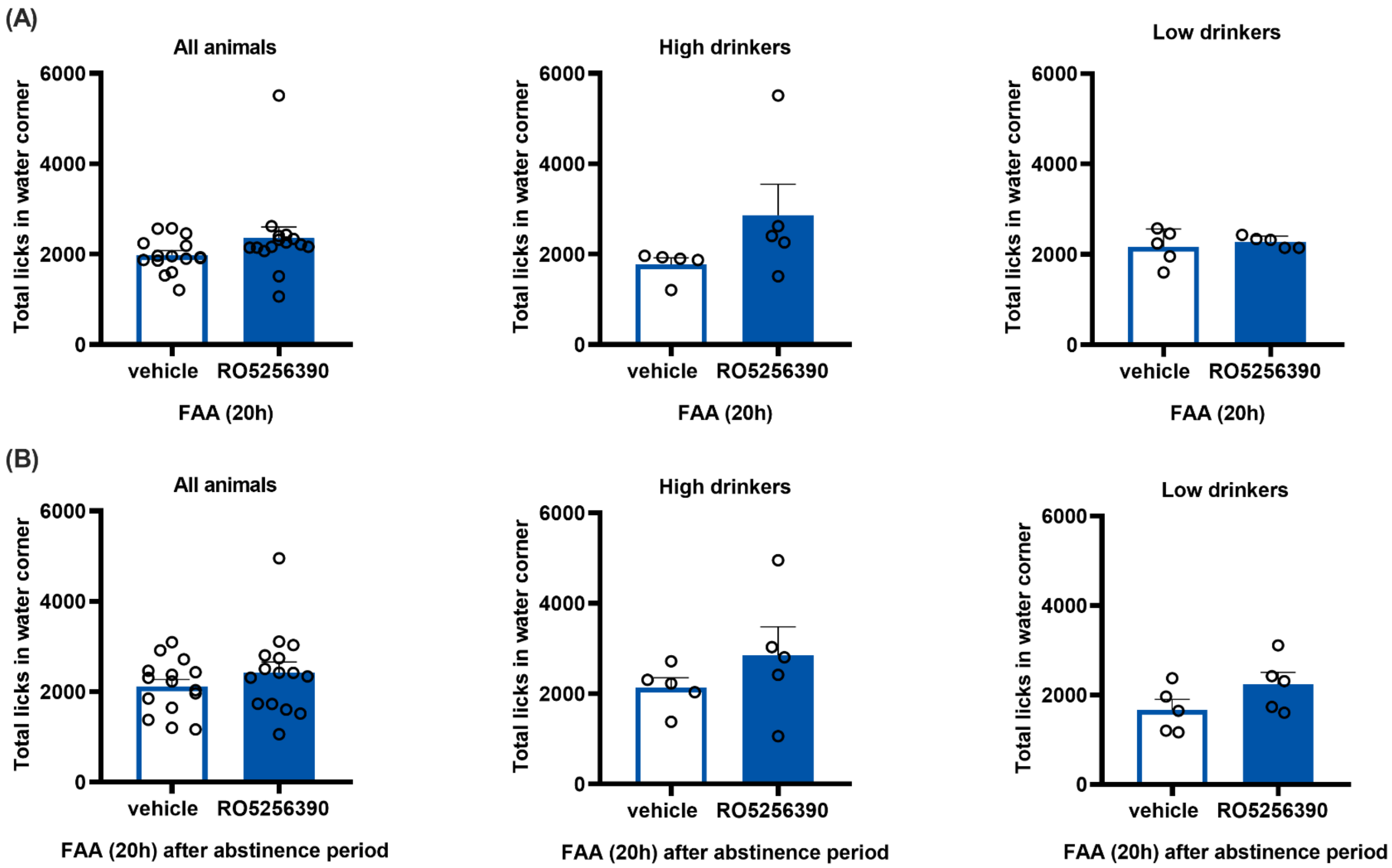
